# Distinct antibody-mediated selection in a narrow-source HIV-1 outbreak among Chinese former plasma donors

**DOI:** 10.1101/167510

**Authors:** Sophie M. Andrews, Yonghong Zhang, Tao Dong, Sarah L. Rowland-Jones, Sunetra Gupta, Joakim ESBJÖRNSSON

## Abstract

The HIV-1 envelope protein mutates rapidly to evade recognition and killing, and is a major target of the humoral immune response and in vaccine development. Identification of common antibody epitopes for vaccine development have been complicated by large variations on both virus (different infecting founder strains) and host genetic levels. We studied HIV-1 envelope *gp120* evolution in 12 Chinese former plasma donors infected with a purportedly single founder virus, with the aim of identifying common antibody epitopes under immune selection. We found five amino acid sites to be under significant positive selection in ≥50% of the patients, and 22 sites housing mutations consistent with antibody-mediated selection. Despite strong selection pressure, some sites housed a limited repertoire of amino acids. Structural modelling revealed that most sites were located on the exposed distal edge of the Gp120 trimer, whilst wholly invariant sites clustered within the centre of the protein complex. Four sites, flanking the V3 hypervariable loop of the Gp120, represent novel antibody epitopes that may be suitable as vaccine candidates.

## INTRODUCTION

The human immunodeficiency virus type 1 (HIV-1) glycoprotein Gp120 is a 120 kDa surface-expressed protein that is essential for viral entry into the cell. It is encoded by the *env* gene, and consists of five variable regions (V1-V5) interspersed between five conserved regions (C1-C5)^1^. The Gp120 forms heterodimers with Gp41 which themselves trimerise, studding the viral membrane at a density of around fourteen copies per virion^2^. Whilst the cellular immune response against HIV-1 targets epitopes dispersed throughout the viral genome, the accessibility of Gp120 on the cell surface makes it the major target of humoral responses.

The humoral response against HIV-1 Gp120 develops rapidly within around four weeks of detectable plasma viral loads^3^, but neutralising antibodies (NAbs) typically only develop after several months of infection^4^. Around two hundred antibodies have been described that recognise the Gp120 protein (LANL Immunology Database; *http://www.hiv.lanl.gov/content/immunology*), and many of the epitopes cluster within the V3 loop. However, the interplay between Gp120 and the adaptive immune response is complex, and the role that antibodies play in the control of infection is a contentious issue. The loss of neutralising activity has been associated with faster disease progression in some individuals^5^. In addition, studies in macaques have indicated that B lymphocyte depletion-associated reductions in NAb titre inversely correlate with viral load, suggesting that the humoral response may contribute at least in part to the control of viral replication^6,7^. However, whilst NAbs do exert selection pressure on the virus^8,9^, the breadth of response does not correlate with or predict progression to AIDS^5,10,11^.

It is commonly believed that the reason why antibody responses may play a limited role in the control of HIV-1 is because the virus can mutate easily to escape neutralization by these responses: as one antibody is evaded, new antibodies are raised but these are evaded again in an on-going cycle^9,12–14^. This view is supported by the observation that HIV-1 is rarely susceptible to neutralisation by contemporaneous antibodies in early infection^15,16^, whilst the same antibodies are able to effectively neutralise historic virus^9,12,17^. However, in clinically latent infection, viral variants evolve susceptibility to neutralisation by contemporaneous NAbs, or to sera sampled much earlier in infection^18–20^. It is therefore possible that antibody responses do play an important role in controlling HIV-1, at least in the latent phase, with re-emergence of variants occurring periodically as the associated NAb responses fall below a certain threshold but then are restored by stimulation by the variant^21^.

Indeed, several apparent paradoxes in HIV-1 pathogenesis and the genetics of host susceptibility can be resolved by assuming that NAbs play an important role in the control of infection, as shown by a recent modelling study^21^. Non-neutralising responses with Fc-related activities – including antibody-dependent cellular cytotoxicity (ADCC) or antibody-mediated cellular viral inhibition (ADCVI) – directed at epitopes of intermediate variability, may also help maintain chronicity of infection. This is consistent with the findings of studies in rhesus macaques demonstrating that simian immunodeficiency virus isolated during clinically latent infection remains susceptible to ADCVI responses from earlier plasma, despite no detectable contemporaneous, autologous neutralising response^22^. A potential therapeutic approach to preventing disease progression may therefore be to develop vaccines that boost and maintain such partially cross-protective responses.

Here we studied *gp120* evolution in a narrow-source cohort of former plasma donors (FPDs) from Henan Province in China (SM cohort)^23^. The FPDs of Henan were infected with a narrow-source of virus through exposure to contaminated equipment, and transfusion with pooled red blood cells during an illegal paid plasma donation scheme in the mid-1990s^24,25^. Owing to the unusually homogeneous route and source of infection, and the narrow timeframe during which patients were exposed, the infecting founder is relatively conserved. This provided us with the unique opportunity to investigate where and how immunological selection pressure drives mutation within *gp120*. Our aim was not only to comprehensively map where antibodies are able to bind in natural infection, but also to understand the constraints placed on mutation in these regions, thereby testing whether antibody escape is a continuous linear process or one in which a limited number of epitopes are re-cycled over time. The identification of such epitopes of limited variability, which are nonetheless targets of natural immunity, may assist in the development of vaccination strategies that prevent viral escape: for example, by incorporating all possible substitution forms into the vaccine.

## METHODS

### Cohort characteristics and sampling

The SM cohort comprises HIV-1 patients from a small rural community in Henan province, China, as described previously^23^. Between 1993 and 1995, the patients were infected with a narrow-source of subtype B’ virus during an illegal paid plasma donation scheme. Few individuals knew that they were infected until HIV screening programmes were implemented in China in 2004. Patients were then recruited to the cohort and gave informed consent for their samples to be used.

Patients with samples collected longitudinally over the course of HIV-1 infection permit detailed evolutionary analysis and identification of sites under selection. Twelve patients from the SM cohort with multiple samples collected between 2010 and 2014 were available for this study (Table S1). Ethical approval was obtained from Beijing You’an Hospital and the University of Oxford Tropical Ethics Committee (OxTREC).

### HIV-1 *env gp120* sequencing and sequence assembly

Viral RNA was isolated from cryopreserved plasma samples and purified using the QIAamp Viral RNA Extraction Kit (Qiagen) followed by reverse transcription using the SuperScript III Reverse Transcriptase System (Invitrogen). *Env gp120* C2-V5 was amplified by nested touchdown PCR (primers listed in Table S2). Amplified DNA was purified using the MinElute Gel Extraction Kit (Qiagen), and ligated into a pCR4-TOPO sequencing vector using the TOPO TA Cloning Kit for Sequencing (Invitrogen). Chemically competent One Shot MAX Efficiency DH5α *E. coli* (Invitrogen) were transformed with the prepared plasmids, and cultured overnight at 37°C. Eighteen colonies were selected for colony PCR (M13F and M13R primers, Table S2), and the resulting products were purified using ExoSAP-IT (Affymetrix) and sequenced to generate forward and reverse reads (Source BioScience). Contigs were assembled and controlled manually using Geneious v9.0.5^26^ (HXB2 *gp120* positions 661-1455, accession number K03455). Sequences were multiple aligned using MUSCLE^27^, and then manually edited in MEGA v6.06^28^. Sequences with premature stop codons or frame-shifts were excluded to control for intra-patient clustering of sequences.

### Inference of infecting founder strain

A consensus sequence was generated for each patient for each time-point, with an ambiguity threshold of 10%. One sequence was selected per patient to generate a dataset with sequences evenly distributed across the sampling period. The sequences were aligned and codon-stripped to a final alignment length of 759 nucleotides. Patient SM007 was excluded from this analysis because it was not possible to conclusively rule out dual- or superinfection as preliminary data exploration demonstrated that sequences from this patient did not resolve monophyletically.

Bayesian Markov Chain Monte Carlo phylogenetic inference - implemented through BEAST v1.8.2 (*http://beast.bio.ed.ac.uk/beast/*)^29^ - was used to estimate the time to the most recent common ancestor (tMRCA). Divergence time was estimated using the SRD06 model^30^, and an uncorrelated lognormal relaxed molecular clock^31^ with a rate prior of 0.001 substitutions per site per year. The Markov chain was run for 100,000,000 generations to allow for adequate mixing, with posterior samples extracted every 10,000 generations. A burn-in period of 10% was applied, and convergence of posterior probabilities was assumed once the effective sample size (ESS) of each parameter exceeded 200, as determined in Tracer v.1.5 (*http://tree.bio.ed.ac.uk/software/tracer/*). Three runs were combined in LogCombiner v1.8.2 (*http://beast.bio.ed.ac.uk/LogCombiner/*) and the mean root height of the tree was calculated. Phylogenetic trees were annotated in FigTree v.1.4.2 (*http://beast.bio.ed.ac.uk/figtree*).

As the SM cohort patients were infected with a narrow source of virus, the sequence of the MRCA was used as a surrogate for the infecting founder. The sequence of the reconstructed ancestor at the tree root from each run following burn-in was extracted, and a consensus sequence was generated from an alignment of these sequences (26,000 sequences) with an ambiguity threshold of 10%.

### Viral subtyping

An unambiguous consensus sequence was generated from the sequences of each time-point for each patient. The sequences were then aligned to the LANL 2005 *gp120* reference dataset (*http://www.hiv.lanl.gov/*), and viral subtyping was performed by Bayesian phylogenetic inference (BEAST v1.8.2). The generalised time-reversible (GTR) nucleotide substitution model plus invariant sites and gamma-distributed rate heterogeneity (GTR+I+G) was used, with a constant size coalescent tree prior, estimated base frequencies, and a strict molecular clock. Following exclusion of burn-in, TreeAnnotator v1.8.2 was used to determine the maximum clade credibility tree (MCC).

### Selection pressure in *env gp120*

The ratio of non-synonymous (dN) to synonymous (dS) mutations was estimated for each codon in patient-specific alignments. Renaissance counting^32,33^ was implemented through BEAST v1.8.2, and the HKY85 nucleotide substitution model^34^, three-site codon partitioning, a strict molecular clock with tip-dating of time stamped sequences were applied. Significant selection was defined as a 95% higher posterior density (HPD) range that did not encompass 1. An alignment representing all patients was then constructed from these data, and the dN/dS estimates were combined across each aligned position. The proportion of patients with virus showing evidence of significant selection pressure in each cohort was calculated.

### Variant characterisation

A variant was defined as any amino acid in any position in the alignment that differed from that present in the inferred founder. Owing to the extensive degree of variation, the hypervariable loops were conservatively stripped from the alignment prior to analysis (final length 179 amino acids). Major variants were defined as variants found in greater than 15% of the sequences. Whilst major variants are canonically defined as those present at a frequency greater than 5%, this value was conservatively tripled as the amplicon was approximately three times more variable than the full-length HIV-1 genome^35^.

### Structural modelling

Homology modelling was implemented through SWISS-MODEL^36–39^ to map the translated SM cohort Gp120 consensus sequence to a cryo-electron microscopy (cryo-EM) crystal structure of a glycosylated HIV-1 envelope trimer (RCSB PDB 3J5M)^40^. The sites of interest were annotated on the modelled structure in PyMOL v1.7.4 (Schrödinger, LLC.).

### Data analysis

Epitope mapping, selection mapping, variant characterisation, biophysical properties, statistical analysis, and plotting were all performed in R (v3.2.3)^41^ via the RStudio (v.0.98.1103) integrated development environment (*http://www.rstudio.com/*). The following libraries were used: dplyr^42^; scales^43^; gridExtra^44^; ggplot2^45^; reshape2^46^. All scripts are available from the authors on request.

## RESULTS

### Cohort characteristics

HIV-1 *env gp120* sequences were recovered from 12 patients in the SM cohort (seven female and five male). The HLA types of those were representative of cohort frequencies^23^. Sequence recovery was 74% from the available specimens, with ten of the patients yielding sequences from two or more time points (Fig. S1A). PCR success was associated with the viral load of the sample. The final dataset of 575 sequences is available from Genbank under accession numbers MF078678-MF079252.

Across the specimens sampled, median viral load was 7,388 copies ml^-1^ of plasma (interquartile range (IQR): 1,612-30,403); median absolute CD4^+^ lymphocyte count was 337 cells µl^-1^ (IQR: 248-400); and median CD4 percentage of lymphocytes was 24% (IQR: 15-30). Demographic and clinical characteristics of the patients sampled are shown in Table S1.

Maximum likelihood phylogenetic analysis showed a star-like relationship between the sequences, with short internal branches between patients, consistent with a narrow-source outbreak (Fig. S1B). Subtype analysis showed that all patient sequences clustered with the CRF15_01B strain: a circulating recombinant virus initially reported in Thailand composed of CRF01_AE with the majority of envelope being subtype B (Figure S1C). Bayesian phylogenetic analysis placed the origin of the SM cohort cluster at around January 1999 (95% HPD October 1988 to November 2006).

### Five positions in Gp120 were under significant positive selection pressure in at least half of the patients irrespective of HLA profile

The median evolutionary rate ratio of positions 1+2 to position 3 among the participants was 0.861 (IQR: 0.779-0.892), indicating an overall purifying selection. The overall ratio was <1 in all patients except SM021 (1.306). Next, we evaluated the intrapatient dN/dS ratio of each codon by renaissance counting^33^. The majority of codons in Gp120 were under either significant positive or negative selection in seven of the ten patients with longitudinal samples (Fig. 1A), and neutral evolution was comparatively rare. Consistent with the 1+2:3 codon rate ratios, only one patient (SM021) appeared to have more residues under significant positive than negative selection pressure.

**Figure 1.**
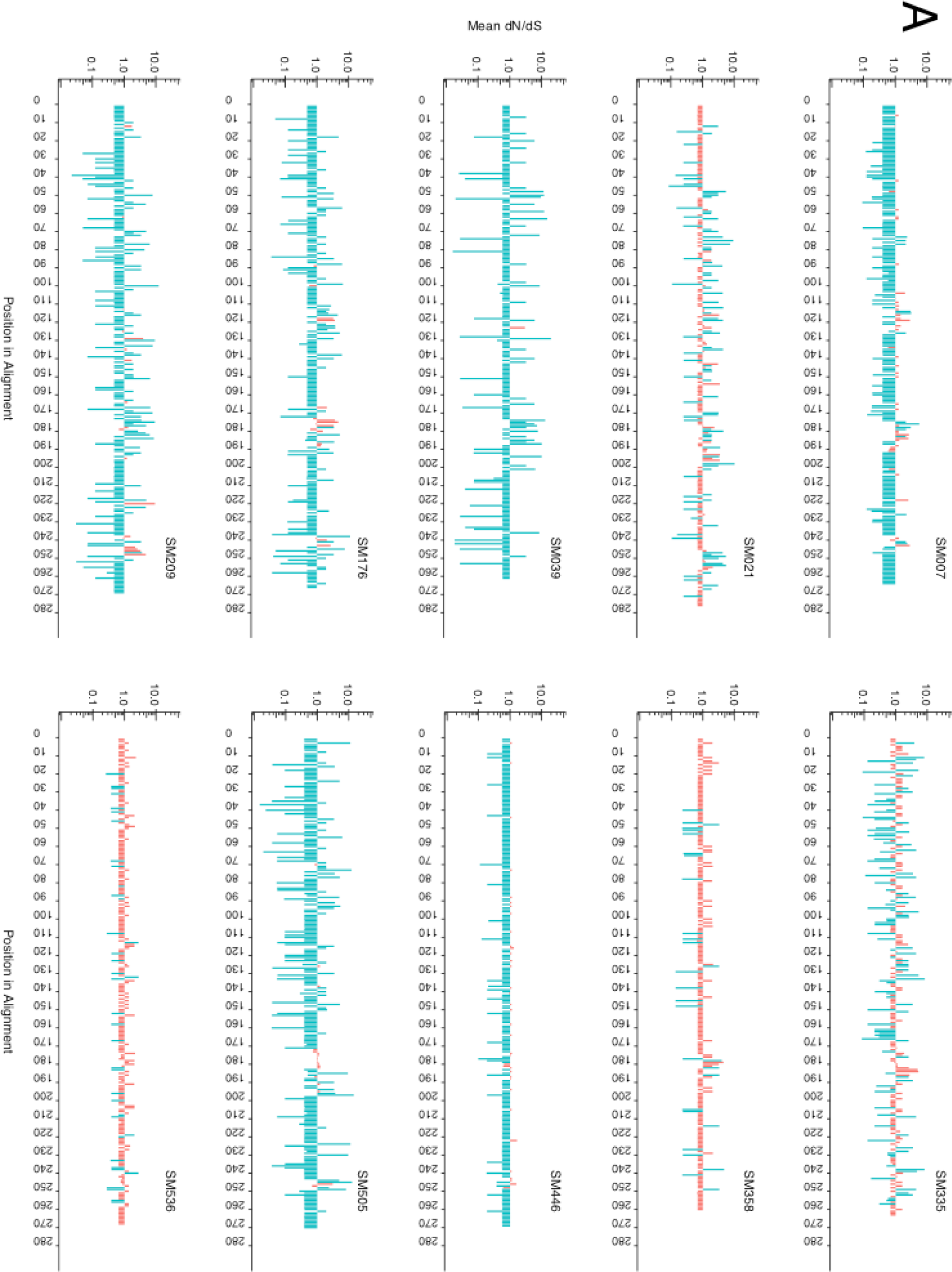
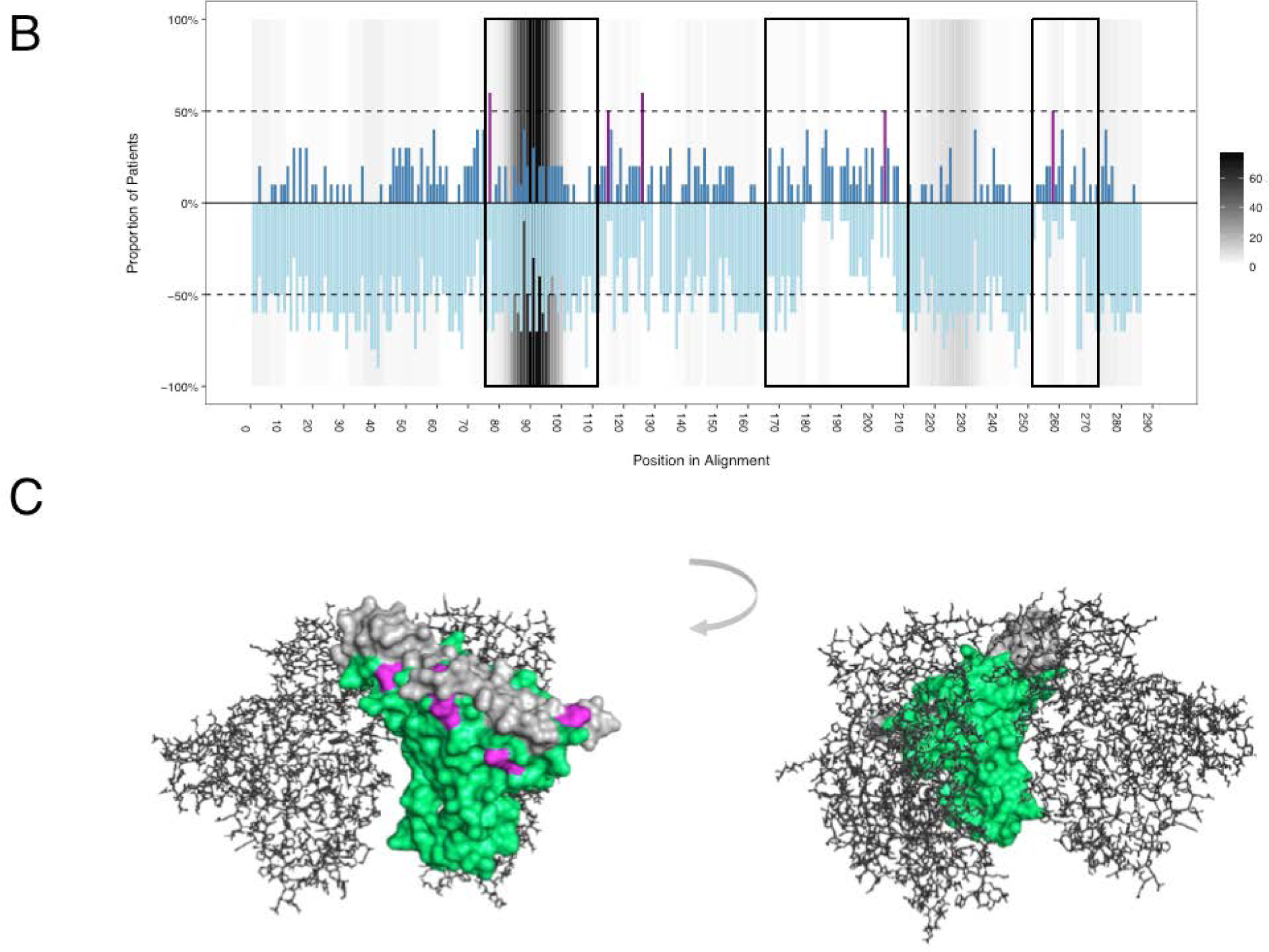
Selection pressure within the HIV-1 envelope Gp120 protein. **A)** Mean dN/dS ratios for each codon within each patient. A dN/dS estimate greater than 1 indicates positive selection whilst an estimate of less than 1 indicates negative selection. Sites under significant selection shown in blue whilst sites that have not reached statistical significance are shown in red and are assumed neutral. Differences in the number of codons across patients are the result of length variation in the V4 and V5 hypervariable loops. **B)** The proportion (absolute) of patients showing evidence of significant positive or negative selection across each aligned codon of the amplicon. Positive selection is shown in dark blue, whilst negative selection is shown in light blue. Sites under significant positive selection in at least 50% of patients are shown in purple. Variable loops V3, V4 and V5 are contained within the three boxes, respectively. The dotted lines denote 50% of patients. Antibody epitope clustering is shown in grey, whereby intensity denotes number of epitopes spanning that residue as reported in the LANL Immunology Database (*http://www.hiv.lanl.gov/content/immunology*). Sequences have been aligned to the HXB2 Gp120 reference sequence (accession number K03455), and position is relative to this alignment. **C)** Homology-modelled structure of the SM cohort consensus Gp120 sequence in surface representation. Variable loops V3, V4 and V5 are shown in grey and sites under significant positive selection in 50% or more patients are shown in purple. Structure has been modelled on a glycosylated HIV-1 Gp120 trimer (RCSB PDB 3J5M)^40^. For clarity, two molecules in the trimer are shown in line representation in grey.

Within the variable loops, substantial negative selection could be seen in V3, but not in V4 and V5 (Fig. 1B). The negative selection in V3 corresponded with a marked increase in the density of neutralising antibody epitopes. Five codons, corresponding to positions T297, A337, S348, D415 and S468 in the HXB2 Gp120 (accession number K03455), were under significant positive selection in 50% or more of the patients irrespective of HLA profile (Figure 1A and Table S1). These sites were further mapped to a homology-modelled structure of the SM cohort Gp120 consensus, and clustered either within or immediately flanking the variable loops on the distal exposed edge of the protein complex (Fig. 1C).

### Significant selection pressure was exerted by the humoral response

We next considered how the virus evolves in response to significant positive selection pressure. For each position in the partial Gp120 sequence, deviations from the inferred founder were detected and 288 unique amino acid variants were recorded across 128 variable sites (Fig. 2A). The remaining 51 positions were completely invariant (29%). Of the variants identified, many were present in a single or small collection of sequences and are likely of limited biological relevance. Major variants were therefore resolved, and to prevent overrepresentation by particular patients, a single time-point was selected for each. Thirty-one major variants were detected in total, across 29 sites (Fig. 2A).

**Figure 2.**
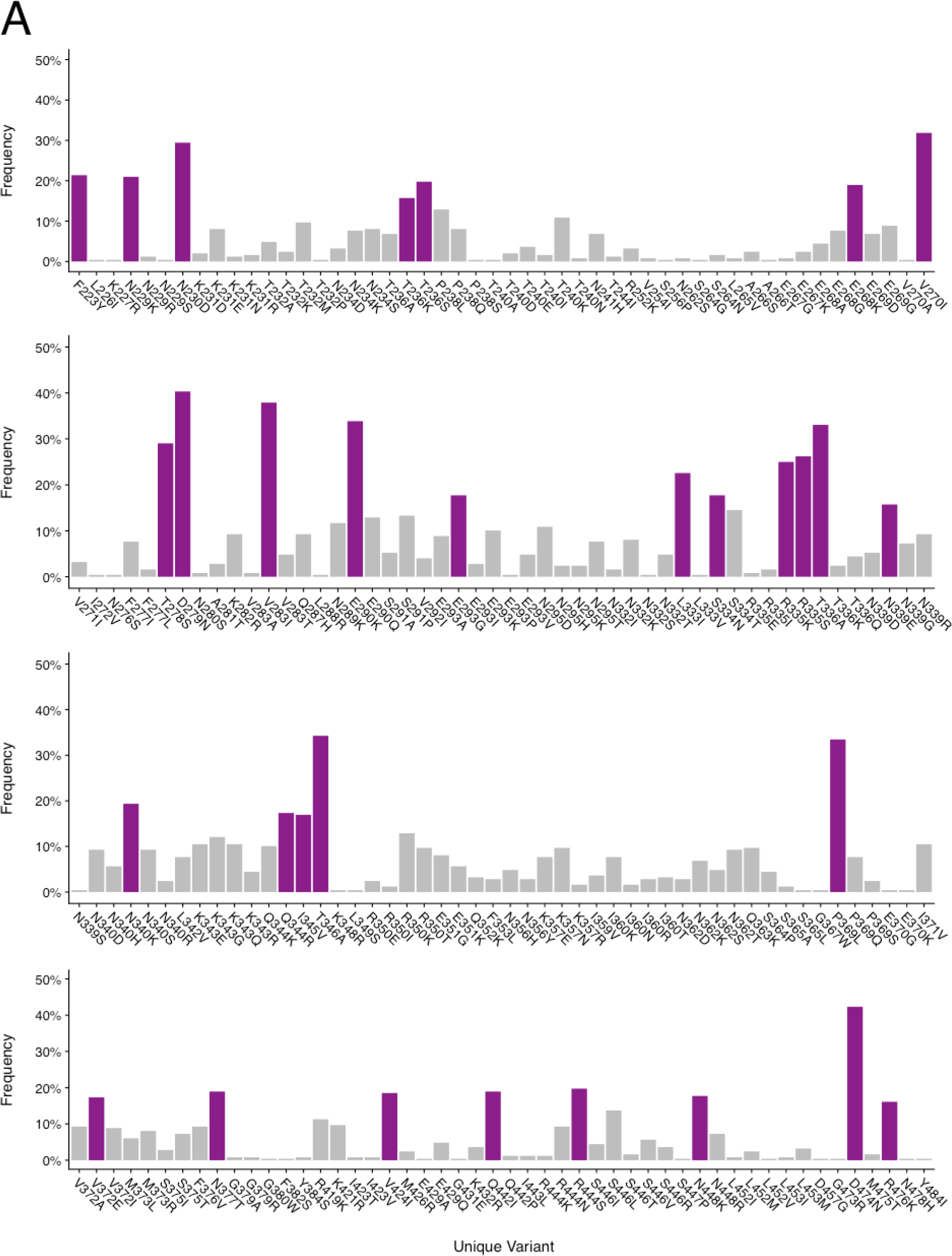
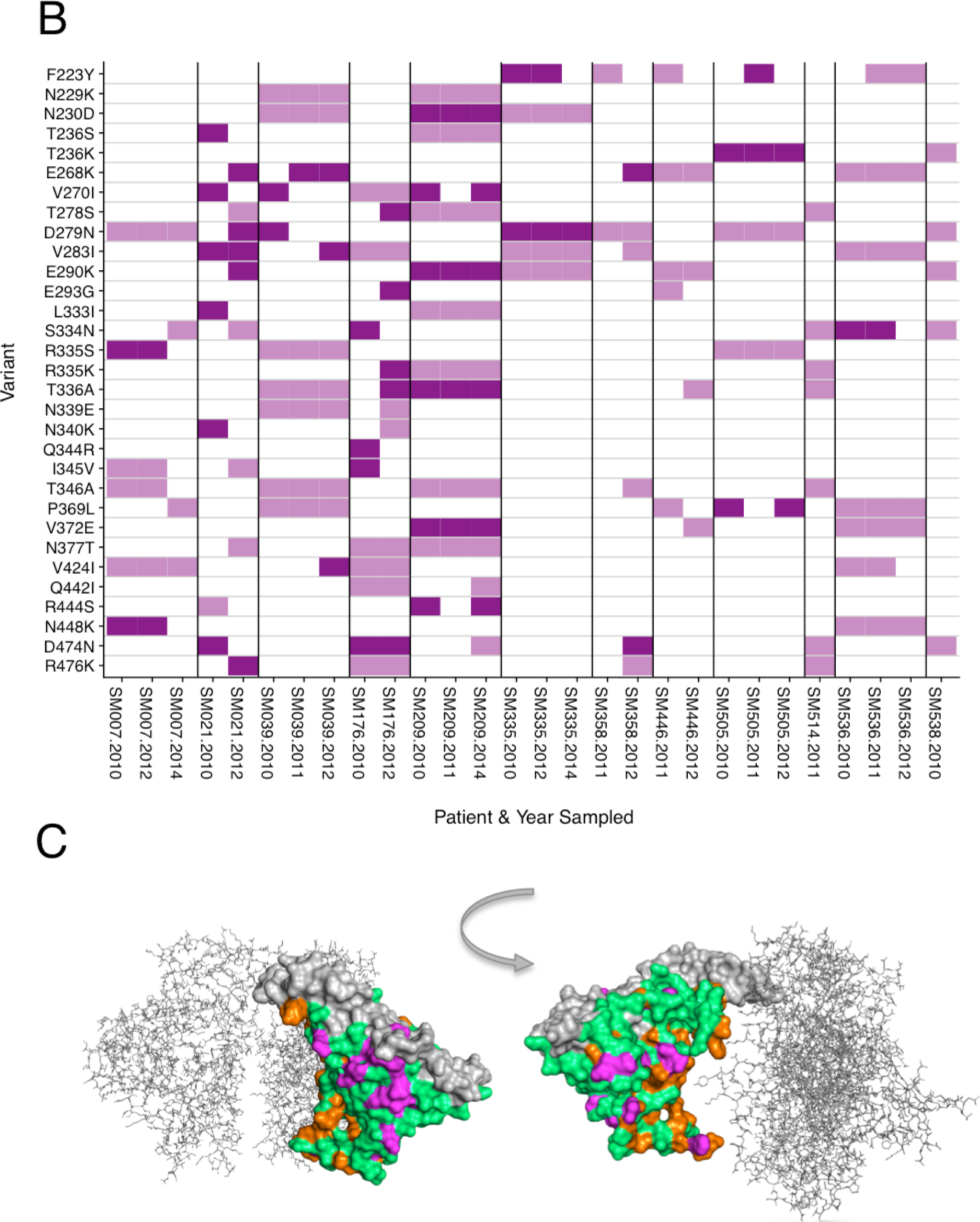
Variants mapping. **A)** Unique variants identified in the SM cohort envelope Gp120 sequences. Major variants present in greater than 15% of the sequences are shown in purple whilst minor variants (<15%) are shown in grey. Position is relative to HXB2 Gp120 (accession number K03455). **B)** Presence of major variants in individual patients of the SM cohort at each time point sampled. Those shown in dark purple are also under significant positive selection pressure within the specified individual, whilst those in light purple are either under significant negative selection or under no significant selection pressure. **C)** Homology modelled structure of the SM cohort consensus Gp120 sequence, showing the variants under significant positive selection in 50% or more of the patients in purple, and all invariant sites in orange. All variants except positions 345 and 424 are visible on the surface of the protein. Variable loops V3, V4 and V5 are shown in grey. Structure has been modelled on a glycosylated HIV-1 Gp120 trimer (RCSB PDB 3J5M)40. For clarity, two molecules in the trimer are shown in line representation in grey.

In patients with longitudinally-sampled sequences, the presence or absence of each major variant was recorded at each time point. Whether these sites were under significant positive selection pressure in that patient was also determined (Fig. 2B). Twenty-four of the 29 sites housing major variants were under significant positive selection pressure in at least one patient, and of these sites, a higher proportion exhibited fluctuating patterns of variant emergence than a consistent pattern wherein the variant was present at all time points sampled (*p*<0.01, two-tailed Fisher’s Exact Test).

Mapping these 24 sites to the homology-modelled structure of the SM cohort Gp120 consensus revealed that all but two were visible on the surface of the protein and likely accessible to antibodies (Fig. 2C). The exceptions were positions 345 and 424. Position 345 houses the major variant I345V, and is contained within the location of a known HLA-A11-restricted CTL epitope^47^. This position was found only to be under significant positive selection pressure in patient SM176, who also expresses HLA-A11. Position 424 houses the CD4 binding site. The wholly invariant sites were similarly mapped, and the overwhelming majority clustered on the inner face of the quaternary structure.

### Despite significant antibody-mediated selection pressure, some sites housed a limited repertoire of amino acids

We next considered the biophysical diversity of amino acids in each of the 22 sites of interest (Fig. 3). In all positions, the biophysical properties of the amino acid in the inferred founder were preserved in approximately 50% or more of the sequences. Little divergence was seen in sites where the inferred founder residue was hydrophobic, with other properties being somewhat more variable. Tabulating the amino acids present in each site also reveals that seven positions flick back and forwards between just two or three amino acids.

**Figure 3.**
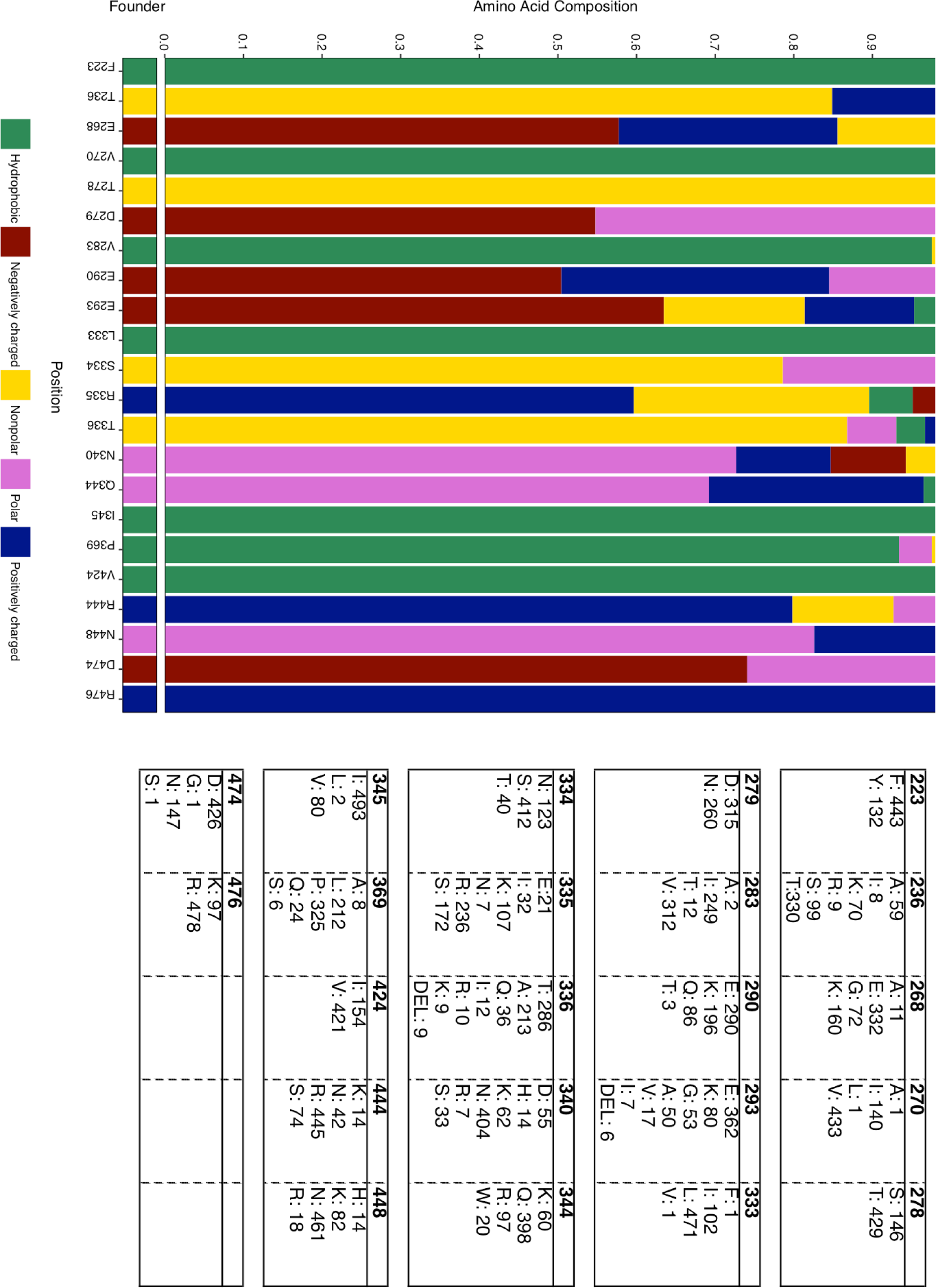
Biophysical properties of amino acids found in sites housing major variants. The biophysical properties of the amino acids found in each site, coloured according to the Lesk format^65^. The table show the composition of each position as the total number of each amino acid found at that site.

### Four novel sites were identified and consistent with antibody activity

The 22 sites were also cross-checked for existing antibody epitopes in the LANL Immunology Database (Table 1). Eighteen of the 22 sites were part of previously described antibody epitopes from different sources (human, mice, or both). In detail, five positions (T236, Q344, I345, V424, and D474) were contained within known human antibody epitopes, whilst 16 positions were contained within epitopes reported in mice. Four sites (V283, E290, L333, S334) flanking the V3 hypervariable loop – which itself houses the majority of antibody epitopes – were identified against which no antibodies have yet been reported. Two of these four sites (V283 and L333) primarily switch between I/V and I/L, respectively.

**Table 1.**
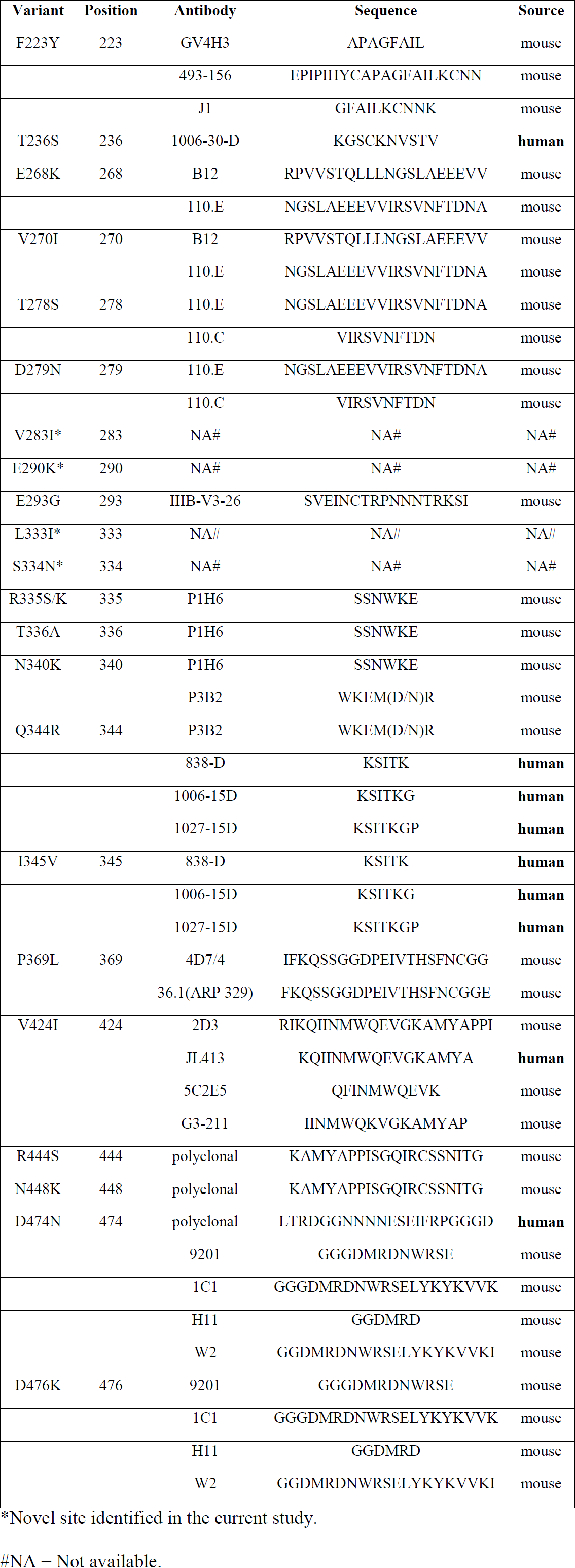
Antibody epitope sequences corresponding to the sites of interest identified in this study. Data made available by the Los Alamos National Laboratory (LANL) Immunology Database (*http://www.hiv.lanl.gov/content/immunology*). Position is relative to HXB2 Gp120 (accession number K03455).

## DISCUSSION

In line with previous observations^48^, mapping the ratio of non-synonymous to synonymous substitutions showed that the majority of sites within the C2-V5 region of Env Gp120 were under negative selection in all but one patient out of ten. Whilst V4 and V5 loops exhibited a dearth of negative selection, the V3 hypervariable loop contained substantial negative selection. Of the five variable loops in Gp120, V3 is the most conserved with amino acid variation restricted to approximately 20% of the loop’s residues^49^. It is also likely that V3 is subject to stronger functional constraints due to its important role in co-receptor binding^50–53^. Moreover, it has been shown that deletion of V3 abrogates viral infectivity^54^.

Several sites within each patient showed evidence of significant positive selection, and five of these were common to at least half of the patients sampled. Structural modelling demonstrated that all but two of the 24 positively selected sites were found on exposed regions of the outer face of the protein complex. CTL epitopes in the HIV-1 Nef protein have been reported to cluster in hydrophobic regions^55^, whilst more recent evidence suggests that their distribution may be random across the genome^56^. Such strong clustering on the surface of the protein is therefore more consistent with antibody-mediated than CTL-mediated selection pressure. The exceptions in terms of surface exposure were positions 345 and 424, which were buried within the protein. Notably, position 345 was found to be under significant positive selection pressure in only one patient, SM176. This position is contained within a known HLA-A11-restricted CTL epitope, which is one of the HLA alleles expressed by patient SM176. It is therefore feasible that this variant has emerged in response to CTL-mediated selection pressure in this patient. Conversely, position 424 is important in CD4 binding^57^, and is contained within a known human antibody epitope. Mutation of this residue to methionine has been shown to increase susceptibility to neutralisation^58^.

We were able to assign most positions to known antibody epitopes in humans and mice. We also identified four novel sites consistent with humoral activity, which are not contained within any known antibody epitopes reported in the literature. These sites flank the V3 hypervariable loop, which is the most epitope dense region of Gp120 (LANL Immunology Database; *https://www.hiv.lanl.gov/content/immunology/*), although this may be due to a bias in reporting stemming from the extensive study of V3 in vaccine design rather than a genuine increase in immune activity. Whilst numerous antibodies against V3 have been described, the cross-neutralisation potential of these antibodies is generally low, reviewed by Hartley *et al*.^59^ Glycosylation, sequence variation, masking by V1-V2, and the specific amino acid make-up of the loop may contribute to this, reviewed by Pantophlet *et al*.^60^. However, some monoclonal and polyclonal antibodies specific to epitopes within V3 have been demonstrated to neutralise diverse HIV-1 strains *in vitro*^61–64^. Two of the new sites exhibit particularly limited amino acid diversity and as such may warrant further investigation as potential components of vaccines targeting shared epitopes of very low diversity within V3.

Indeed, despite evidence for significant antibody-mediated selection pressure, some sites were relatively conserved in terms of their composition, containing only a limited number of biophysically similar amino acids. This is likely due to functional or structural constraints on the protein, and may reduce the ability of the virus to successfully escape antibodies targeting these regions. We also identified sites containing biophysically diverse amino acids that may be contained within epitopes eliciting highly effective antibody responses which cycle continuously between a limited number of biophysically distinct forms throughout chronic infection, as predicted by a previous modelling study^21^. Consistent with this model, we found evidence within individuals that some sites contain major variants that appear and disappear over time, such as proline to leucine in position 369 and arginine to serine in position 444.

In summary, our data provide insight into how and where the surface of Gp120 is mutating over the course of clinically latent infection, and indicate that humoral immunity is the likely driving factor of such changes. Our detailed analysis of HIV-1 evolution and selection within hosts infected with a narrow-source virus allowed us to identify amino acid sites under positive selection that was likely attributed to host factors. Importantly, this comprehensive mapping resulted in the identification of both previously described and novel constrained antibody and T-cell epitopes. These sites are likely crucial to viral envelope function, and may aid in the development of future drugs and vaccines.

## ACKNOWLEDGEMENTS

We would like to thank the participants of the SM cohort who contributed samples towards this study. We would like to acknowledge the following funding bodies: SMA has received funding from the Wellcome Trust (099815). JE has received funding from the Swedish Research Council (350-2012-6628, 2016-01417) and the Swedish Society for Medical Research (SA-2016). SG has received funding from the European Research Council under the European Union’s Seventh Framework Programme (FP7/2007-2013) / ERC grant agreement no. 268904–DIVERSITY. YZ’s work has been supported by the National Natural Science Foundation of China (81271842, 81320108017), Beijing Natural Science Foundation (7132098), Beijing Municipal Administration of Hospitals (XMLX201411), Beijing Key Laboratory (BZ0373), and Capital Health Development (TG-2015-19). Initial sample collection was funded by the Li Ka Shing Foundation.

## AUTHOR CONTRIBUTIONS

S.M.A performed the experiments. S.M.A and J.E analysed the data. S.M.A, J.E, S.R-J, T.D, and S.G. contributed substantially to the conception and design of the study. Y.Z. sourced the samples used in the study. S.M.A wrote the manuscript, S.M.A, J.E, S.G. and S.R-J edited the manuscript, and all authors gave approval of the manuscript for submission.

## DECLARATIONS

None.

## ETHICS APPROVAL AND CONSENT TO PARTICIPATE

Ethical approval was obtained for this study from Beijing You’an Hospital and the University of Oxford Tropical Ethics Committee (OxTREC).

## CONSENT FOR PUBLICATION

Not applicable

## AVAILABILITY OF DATA AND MATERIALS

The datasets generated and analysed during the current study are available in the Genbank repository (accession numbers: MF078678-MF079252). Custom R scripts used in the analysis of these data are available from the authors on request.

## REFERENCES

1 Wyatt, R. et al. The antigenic structure of the HIV gp120 envelope glycoprotein. Nature 393, 705-711 (1998).

2 Zhu, P. et al. Distribution and three-dimensional structure of AIDS virus envelope spikes. Nature 441, 847-852 (2006).

3 Tomaras, G. D. et al. Initial B-cell responses to transmitted human immunodeficiency virus type 1: virion-binding immunoglobulin M (IgM) and IgG antibodies followed by plasma anti-gp41 antibodies with ineffective control of initial viremia. J. Virol. 82, 12449-12463 (2008).

4 Legrand, E. et al. Course of specific T lymphocyte cytotoxicity, plasma and cellular viral loads, and neutralizing antibody titers in 17 recently seroconverted HIV type 1-infected patients. AIDS Res. Hum. Retroviruses 13, 1383-1394 (1997).

5 Cecilia, D., Kleeberger, C., Munoz, A., Giorgi, J. V. & Zolla-Pazner, S. A longitudinal study of neutralizing antibodies and disease progression in HIV-1-infected subjects. J. Infect. Dis. 179, 1365-1374 (1999).

6 Schmitz, J. E. et al. Effect of humoral immune responses on controlling viremia during primary infection of rhesus monkeys with simian immunodeficiency virus. J. Virol. 77, 2165-2173 (2003).

7 Miller, C. J. et al. Antiviral antibodies are necessary for control of simian immunodeficiency virus replication. J. Virol. 81, 5024-5035 (2007).

8 Frost, S. D. et al. Neutralizing antibody responses drive the evolution of human immunodeficiency virus type 1 envelope during recent HIV infection. Proc. Natl. Acad. Sci. USA 102, 18514-18519 (2005).

9 Richman, D. D., Wrin, T., Little, S. J. & Petropoulos, C. J. Rapid evolution of the neutralizing antibody response to HIV type 1 infection. Proc. Natl. Acad. Sci. USA 100, 4144-4149 (2003).

10 Piantadosi, A. et al. Breadth of neutralizing antibody response to human immunodeficiency virus type 1 is affected by factors early in infection but does not influence disease progression. J. Virol. 83, 10269-10274 (2009).

11 Euler, Z. et al. Cross-reactive neutralizing humoral immunity does not protect from HIV type 1 disease progression. J. Infect. Dis. 201, 1045-1053 (2010).

12 Albert, J. et al. Rapid development of isolate-specific neutralizing antibodies after primary HIV-1 infection and consequent emergence of virus variants which resist neutralization by autologous sera. AIDS 4, 107-112 (1990).

13 Moog, C., Fleury, H. J., Pellegrin, I., Kirn, A. & Aubertin, A. M. Autologous and heterologous neutralizing antibody responses following initial seroconversion in human immunodeficiency virus type 1-infected individuals. J. Virol. 71, 3734-3741 (1997).

14 Wei, X. et al. Antibody neutralization and escape by HIV-1. Nature 422, 307-312 (2003).

15 Von Gegerfelt, A., Albert, J., Morfeldt-Manson, L., Broliden, K. & Fenyo, E. M. Isolate-specific neutralizing antibodies in patients with progressive HIV-1-related disease. Virology 185, 162-168 (1991).

16 Bjorling, E. et al. Autologous neutralizing antibodies prevail in HIV-2 but not in HIV-1 infection. Virology 193, 528-530 (1993).

17 Geffin, R., Hutto, C., Andrew, C. & Scott, G. B. A longitudinal assessment of autologous neutralizing antibodies in children perinatally infected with human immunodeficiency virus type 1. Virology 310, 207-215 (2003).

18 Aasa-Chapman, M. M. I. et al. In vivo emergence of HIV-1 highly sensitive to neutralizing antibodies. Plos One 6, doi:10.1371/journal.pone.0023961 (2011).

19 Mahalanabis, M. et al. Continuous viral escape and selection by autologous neutralizing antibodies in drug-naive human immunodeficiency virus controllers. Journal of Virology 83, 662-672, doi:10.1128/jvi.01328-08 (2009).

20 Chaillon, A. et al. Human immunodeficiency virus type-1 (HIV-1) continues to evolve in presence of broadly neutralizing antibodies more than ten years after infection. PLoS One 7, e44163, doi:10.1371/journal.pone.0044163 (2012).

21 Wikramaratna, P. S., Lourenço, J., Klenerman, P., Pybus, O. G. & Gupta, S. Effects of neutralizing antibodies on escape from CD8+ T-cell responses in HIV-1 infection. Philosophical Transactions of the Royal Society B: Biological Sciences 370, 20140290, doi:10.1098/rstb.2014.0290 (2015).

22 Asmal, M. et al. Antibody-Dependent Cell-Mediated Viral Inhibition Emerges after Simian Immunodeficiency Virus SIVmac251 Infection of Rhesus Monkeys Coincident with gp140-Binding Antibodies and Is Effective against Neutralization-Resistant Viruses. Journal of Virology 85, 5465-5475, doi:10.1128/JVI.00313-11 (2011).

23 Dong, T. et al. Extensive HLA-driven viral diversity following a narrow-source HIV-1 outbreak in rural China. Blood 118, 98-106 (2011).

24 Zhang, L. et al. Molecular characterization of human immunodeficiency virus type 1 and hepatitis C virus in paid blood donors and injection drug users in china. J. Virol. 78, 13591-13599 (2004).

25 Kaufman, J. & Jing, J. China and AIDS - the time to act is now. Science 296, 2339-2340 (2002).

26 Kearse, M. et al. Geneious Basic: an integrated and extendable desktop software platform for the organization and analysis of sequence data. Bioinformatics 28, 1647-1649 (2012).

27 Edgar, R. C. MUSCLE: multiple sequence alignment with high accuracy and high throughput. Nucleic Acids Res. 32, 1792-1797 (2004).

28 Tamura, K., Stecher, G., Peterson, D., Filipski, A. & Kumar, S. MEGA6: Molecular Evolutionary Genetics Analysis version 6.0. Mol. Biol. Evol. 30, 2725-2729 (2013).

29 Drummond, A. J. & Rambaut, A. BEAST: Bayesian evolutionary analysis by sampling trees. BMC Evol. Biol. 7, 214 (2007).

30 Shapiro, B., Rambaut, A. & Drummond, A. J. Choosing appropriate substitution models for the phylogenetic analysis of protein-coding sequences. Mol. Biol. Evol. 23, 7-9 (2006).

31 Drummond, A. J., Ho, S., Phillips, M. & Rambaut, A. Relaxed phylogenetics and dating with confidence. PLoS Biol. 4, e88 (2006).

32 O’Brien, J. D., Minin, V. N. & Suchard, M. A. Learning to count: robust estimates for labeled distances between molecular sequences. Mol. Biol. Evol. 26, 801-814 (2009).

33 Lemey, P., Minin, V. N., Bielejec, F., Kosakovsky Pond, S. L. & Suchard, M. A. A counting renaissance: combining stochastic mapping and empirical Bayes to quickly detect amino acid sites under positive selection. Bioinformatics 28, 3248-3256 (2012).

34 Hasegawa, M., Kishino, H. & Yano, T.-a. Dating of the human-ape splitting by a molecular clock of mitochondrial DNA. Journal of Molecular Evolution 22, 160-174, doi:10.1007/bf02101694 (1985).

35 Snoeck, J., Fellay, J., Bartha, I., Douek, D. C. & Telenti, A. Mapping of positive selection sites in the HIV-1 genome in the context of RNA and protein structural constraints. Retrovirology 8, 1-8 (2011).

36 Arnold, K., Bordoli, L., Kopp, J. & Schwede, T. The SWISS-MODEL workspace: a web-based environment for protein structure homology modelling. Bioinformatics 22, 195-201 (2006).

37 Kiefer, F., Arnold, K., Kunzli, M., Bordoli, L. & Schwede, T. The SWISS-MODEL Repository and associated resources. Nucleic Acids Res. 37, D387-392 (2009).

38 Biasini, M. et al. SWISS-MODEL: modelling protein tertiary and quaternary structure using evolutionary information. Nucleic Acids Res. 42, W252-258 (2014).

39 Guex, N., Peitsch, M. C. & Schwede, T. Automated comparative protein structure modeling with SWISS-MODEL and Swiss-PdbViewer: a historical perspective. Electrophoresis 30 **Suppl** 1, S162-173 (2009).

40 Lyumkis, D. et al. Cryo-EM structure of a fully glycosylated soluble cleaved HIV-1 envelope trimer. Science 342, 1484-1490 (2013).

41 R Core Team. (Vienna, Austria, 2015).

42 Wickham, H. & Francois, R. (2015).

43 Wickham, H. (2016).

44 Auguie, B. (2016).

45 Wickham, H. ggplot2: Elegant Graphics for Data Analysis. (Springer-Verlag New York, 2009).

46 Wickham, H. Reshaping Data with the reshape Package. Journal of Statistical Software 21, 1-20 (2007).

47 Sriwanthana, B. et al. HIV-specific cytotoxic T lymphocytes, HLA-A11, and chemokine-related factors may act synergistically to determine HIV resistance in CCR5 delta32-negative female sex workers in Chiang Rai, northern Thailand. AIDS Res Hum Retroviruses 17, 719-734, doi:10.1089/088922201750236997 (2001).

48 Edwards, C. T. T. et al. Evolution of the human immunodeficiency virus envelope gene is dominated by purifying selection. Genetics 174, 1441-1453 (2006).

49 Zolla-Pazner, S. & Cardozo, T. Structure–function relationships of HIV-1 envelope sequence-variable regions provide a paradigm for vaccine design. Nature reviews. Immunology 10, 527-535 (2010).

50 Cormier, E. G. & Dragic, T. The crown and stem of the V3 loop play distinct roles in human immunodeficiency virus type 1 envelope glycoprotein interactions with the CCR5 coreceptor. J. Virol. 76, 8953-8957 (2002).

51 Shioda, T., Levy, J. A. & Cheng-Mayer, C. Small amino acid changes in the V3 hypervariable region of gp120 can affect the T-cell-line and macrophage tropism of human immunodeficiency virus type 1. Proc. Natl. Acad. Sci. USA 89, 9434-9438 (1992).

52 Cardozo, T. et al. Structural basis for coreceptor selectivity by the HIV type 1 V3 loop. AIDS Res. Hum. Retroviruses 23, 415-426 (2007).

53 Fenyo, E. M., Esbjornsson, J., Medstrand, P. & Jansson, M. Human immunodeficiency virus type 1 biological variation and coreceptor use: from concept to clinical significance. Journal of internal medicine 270, 520-531, doi:10.1111/j.1365-2796.2011.02455.x (2011).

54 Cao, J. et al. Replication and neutralization of human immunodeficiency virus type 1 lacking the V1 and V2 variable loops of the gp120 envelope glycoprotein. J. Virol. 71, 9808-9812 (1997).

55 Lucchiari-Hartz, M. et al. Differential proteasomal processing of hydrophobic and hydrophilic protein regions: contribution to cytotoxic T lymphocyte epitope clustering in HIV-1-Nef. Proc. Natl. Acad. Sci. USA 100, 7755-7760 (2003).

56 Schmid, B. V., Keşmir, C. & de Boer, R. J. The distribution of CTL epitopes in HIV-1 appears to be random, and similar to that of other proteomes. BMC Evol. Biol. 9, 1-15 (2009).

57 Kwong, P. D. et al. Structure of an HIV gp120 envelope glycoprotein in complex with the CD4 receptor and a neutralizing human antibody. Nature 393, 648-659 (1998).

58 Ringe, R. et al. A single amino acid substitution in the C4 region in gp120 confers enhanced neutralization of HIV-1 by modulating CD4 binding sites and V3 loop. Virology 418, 123-132 (2011).

59 Hartley, O., Klasse, P. J., Sattentau, Q. J. & Moore, J. P. V3: HIV’s switch-hitter. AIDS Res. Hum. Retroviruses 21, 171-189 (2005).

60 Pantophlet, R., Wrin, T., Cavacini, L. A., Robinson, J. E. & Burton, D. R. Neutralizing activity of antibodies to the V3 loop region of HIV-1 gp120 relative to their epitope fine specificity. Virology 381, 251-260 (2008).

61 Conley, A. J. et al. Neutralization of primary human immunodeficiency virus type 1 isolates by the broadly reactive anti-V3 monoclonal antibody, 447-52D. J. Virol. 68, 6994-7000 (1994).

62 Gorny, M. K. et al. Cross-clade neutralizing activity of human anti-V3 monoclonal antibodies derived from the cells of individuals infected with non-B clades of human immunodeficiency virus type 1. J. Virol. 80, 6865-6872 (2006).

63 Hioe, C. E. et al. Anti-V3 monoclonal antibodies display broad neutralizing activities against multiple HIV-1 subtypes. PLoS One 5, e10254 (2010).

64 Corti, D. et al. Analysis of memory B cell responses and isolation of novel monoclonal antibodies with neutralizing breadth from HIV-1-infected individuals. PLoS One 5, e8805 (2010).

65 Lesk, A. M. Introduction to Bioinformatics. (Oxford University Press, 2002).

